# MyoBack: A Musculoskeletal Model of the Human Back with Integrated Exoskeleton

**DOI:** 10.1101/2025.03.13.643057

**Authors:** Rohan Walia, Morgane Billot, Kevin Garzon-Aguirre, Swathika Subramanian, Huiyi Wang, Mohamed Irfan Refai, Guillaume Durandau

**Author notes:** These authors contributed equally to this manuscript (Rohan Walia, Morgane Billot, Kevin Garzon-Aguirre).

## Abstract

Given the challenges of real-life experimentation, musculoskeletal simulation models could become essential in biomedical research. This is especially critical for the human back, a key structure involved in daily movements, where modeling and simulation could streamline design and support the development of treatments and robotic rehabilitation techniques, such as exoskeletons. However, musculoskeletal simulation engines are computationally demanding and lack contact dynamics, restricting current models’ use in studying prolonged behaviors or optimizing system design while maintaining physiological accuracy. To overcome this limitation, this work proposes MyoBack, a human back model part of the MyoSuite framework relying on the physics engine MuJoCo. This model is derived from a physiologically accurate model built in the state-of-the-art musculoskeletal simulation software OpenSim and replicates the latter’s kinematic properties accurately, with some discrepancies regarding muscle dynamics stemming from engine differences. The MyoBack model was also validated empirically by integrating a passive back exoskeleton in simulation and comparing forces exerted on the back with values from experimental trials. Over different tasks, the model reproduced measured force progressions well, resulting in RMSE = 11% for a stoop and RMSE = 16% for a squat motion pattern relative to peak forces. The MyoBack model can be accessed here: https://github.com/rohwalia/MyoBack

## I. INTRODUCTION

Human biomechanics is a rapidly growing field that attempts to understand the intricate interplay between biology and mechanics in human movement, structure, and function. In particular, modeling the back presents significant challenges due to its complex anatomy and the delicate balance required to maintain posture and mobility. The back contributes to everyday movements such as running or lifting objects [1], [2], [3], [4]. Exactly these kinds of common tasks are affected by musculoskeletal disorders (MSDs) prevalent in around 20% of the global population [5], much of it work-related [6], [7] and occurring primarily in the lower back [8]. These issues, in turn, can lead to other complications such as gait disorders, particularly in the elderly [9], limiting quality of life notably.

To alleviate these problems, assistance systems such as exoskeletons are becoming increasingly widespread and hold significant potential to enhance human mobility [10]. However, their development is human resource-intensive and time-consuming [11], [12], [10]. Integrating design and testing environments through modeling and simulation can significantly enhance efficiency, reducing time and hardware costs [13], [14]. Simulation also enables diverse and repeatable testing for optimal system performance. Hence, reliable human body models are essential to better understand musculoskeletal dynamics, identify the causes of MSDs, assess risks of repetitive or prolonged actions, and aid in the customization of medical devices for patient-specific treatment solutions.

Currently, a wide variety of human models are available. For instance, a relatively simple representation of the human body is the Human Dynamic Model [15] which simulates movements based on motion capture data. This model incorporates sufficient degrees of freedom to reproduce complex movements and approximate mechanical energy expenditure during tasks. On the other hand, there are physics-based musculoskeletal simulation engines such as OpenSim [16], AnyBody [17], and SIMM [18] which focus on making detailed and physiologically accurate models of the musculoskeletal system. These tools are extensively used in human neuromechanical control, human-robot interaction, and rehabilitation. Several of these human body models already include the back and have been used in a variety of research projects, for example, to study the effect of unilateral knee extension restriction on the back during gait [19] or to analyze the biomechanical effects of different bag-carrying styles on the back region [20], among others [21], [22], [23].

These musculoskeletal simulation engines are computationally intensive and offer limited support for complex contact interactions. Consequently, most current human body models are impractical for long-term simulations or highfrequency task repetitions while maintaining physiological accuracy. As a result, they are unsuitable for studying longterm behavior or for optimization purposes in system design. Notably, the proprietary software Hyfydy [24] provides a valid approach to this problem. Still, since it is not opensource, using it for custom use cases (e.g., testing personal exoskeleton) in different settings, such as research and industrial, might be difficult. To meet the needs of physiological accuracy, contact dynamics, speed, and accessibility, the use of MuJoCo (Multi-Joint dynamics with Contact) [25] is particularly relevant. MuJoCo is a state-of-the-art open-source physics engine designed for model-based control, offering efficient computational capabilities for multi-joint dynamics and contact mechanics. Importantly, it has the capacity to simulate muscle dynamics while supporting contact-rich interactions. In particular, MyoSuite [26], a software framework based on MuJoCo, is more than two orders of magnitude faster than state-of-the-art musculoskeletal models. This advantage is particularly crucial for highly detailed musculoskeletal models, including our model which features a tenfold increase in muscle patterning and joint degrees of freedom complexity with 210 actuators [27]. To date, various models have been implemented (e.g., arm, leg) but not yet a model of the back. Integrating a back model into MyoSuite would, therefore, enable complex human movements involving the back to be simulated quickly and accurately. This has huge potential in the testing and development of exoskeletons and could be beneficial in rehabilitation, ergonomic design, and personalized healthcare solutions.

Therefore, we developed MyoBack, a back model part of the MyoSuite framework based on an existing OpenSim model [28]. Section II details how the MyoBack model is created and validated by comparing the OpenSim and the derived MuJoCo model with respect to kinematics and dynamics. Furthermore, a passive exoskeleton was developed within the simulation to validate the back model against empirical data. Section III explores the impact of the passive exoskeleton on various back-involved movements by integrating it with the back model and comparing the simulation outputs to experimental results.

## II. METHODS

This section first describes the musculoskeletal model conversion tool [27], MyoConverter, that was used to convert the back model from Opensim to MuJoCo. Then, the manual tuning performed with the output of the conversion tool to complete the final MyoSuite back model is explained. Finally, it is outlined how a passive exoskeleton is added to the back model. This is the basis for experimental validation following the setup and protocol developed in section III.

### A. MyoConverter

We derive our MyoSuite back model (.xml) from an OpenSim model (.osim) [28] using MyoConverter. This conversion tool optimizes muscle kinematics and kinetics and has been tested extensively on other musculoskeletal models such as neck, lower limb and arm [27]. The OpenSim model contains 210 muscle-tendon units (MTUs) and 18 joints. The model can be controlled by 3 “virtual” DoFs that map to the real joints: flexion extension, lateral bending and axial rotation. It has also been incorporated into a fullbody lumbar spine model [29] with a general validation regarding model parameters and muscle output. In addition, a slightly adapted model [30] was explicitly validated for lifting motions, demonstrating its suitability to evaluate changes in lumbar loading during lifting. Given its extensive validation, particularly in terms of muscle activation and joint moments, the OpenSim model is regarded as the state-of-the-art reference, and the goal of MyoConverter is to align the MuJoCo model to it as closely as possible.

The conversion tool consists of three steps:

- Step 1 – Geometry Conversion: parse multi-body kinematic tree containing the fundamental elements of musculoskeletal models (body segments, joints, muscles, wrapping bodies, etc.) from OpenSim into MuJoCo.
- Step 2 – Wrapping optimization: optimize wrapping constraints to guarantee that the muscle pathways in MuJoCo are physiologically reasonable like in OpenSim while accounting for the fact that MuJoCo’s wrapping surfaces are simple geometric objects (e.g., spheres, cylinders), which prevents an exact replication of Open-Sim muscle pathways but enhances computational efficiency, and check the resulting moment arms exerted by each muscle.
- Step 3 – Force property optimization: optimize muscle force parameters in MuJoCo to approximate the Open-Sim model’s force curves and check the force-length-activation.

The conversion tool needed to be modified to convert the back model. Mainly, the handling of edge cases occurring in the OpenSim model that had not been implemented in MyoConverter, e.g., coupled constraints of the *Piecewise-LinearFunction* type, was added.

### B. MuJoCo: MyoBack

The output model from MyoConverter had a few issues. The wrapping pathways of some muscles around a geometric body for specific joint configurations were discontinuous, leading to discrete jumps in muscle-tendon length (Fig.1a). Also, MyoConverter did not parse some geometries correctly from the OpenSim model (Fig.1c). The mapping of the 3 “virtual” DoFs to the real joints of the model had to be fixed as well (Fig.1b,d) by changing the equality constraints [31].

**Fig. 1:**
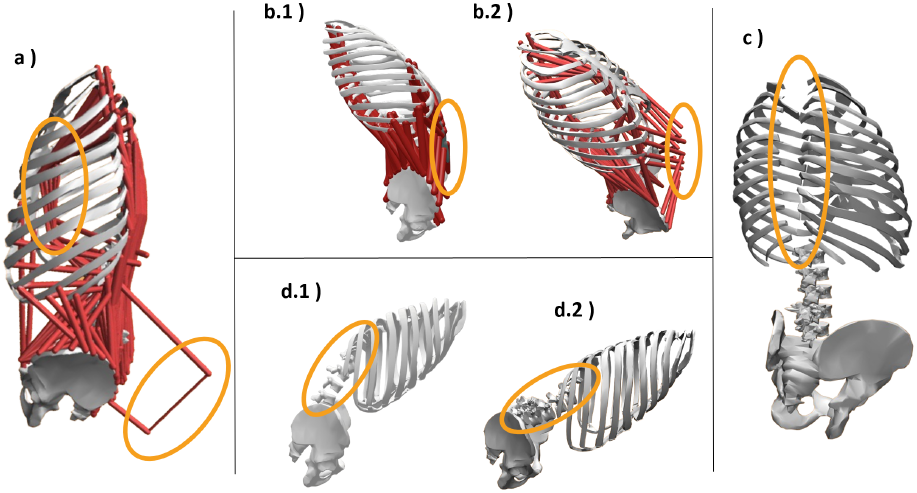
Model problems after conversion a) Jumping muscles b) Immobile abdomen 1) Opensim 2) MuJoCo c) Non-curved spine 1) Opensim 2) MuJoCo d) Absent thoracic vertebrae.

This back model was compared to the OpenSim model with respect to kinematics by observing end point positions (Fig.2a), moment arms exerted by muscles on joints (Fig.2b) and force-length-activation (Fig.2c). The results of this comparison are shown for relevant geometric bodies in the kinematic chain for the former and representative back muscles for the latter two.

**Fig. 2:**
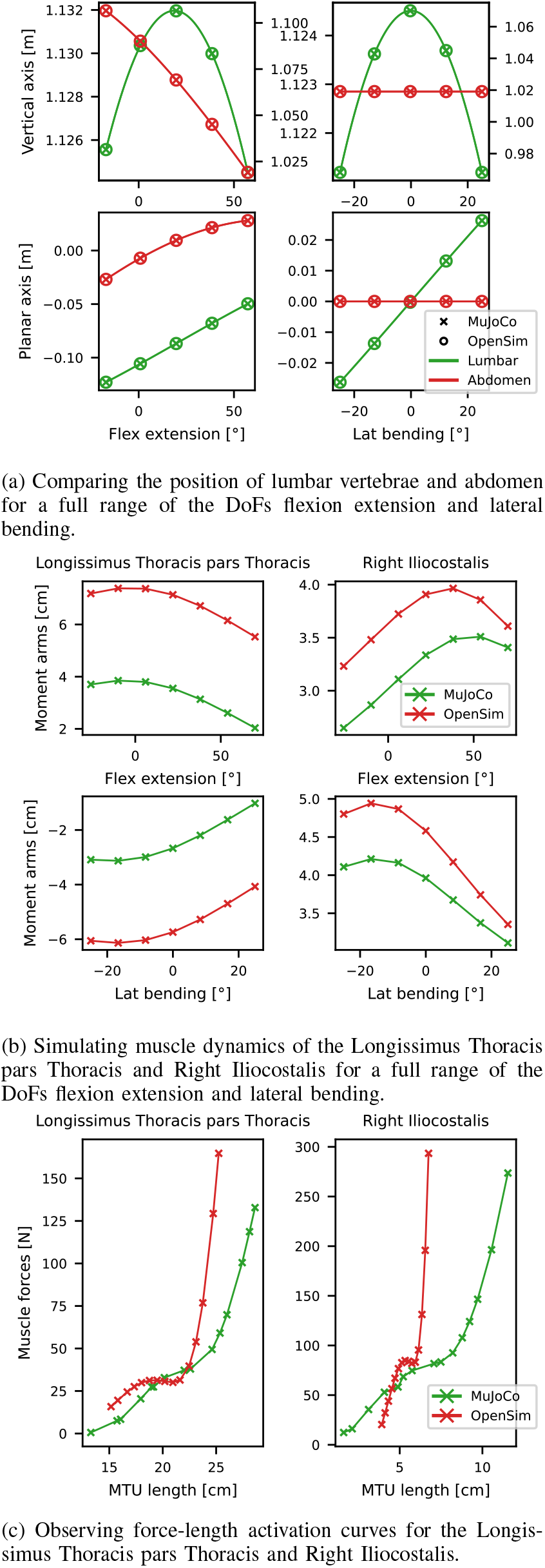
Comparative analysis of the MyoSuite back model and the reference OpenSim model.

### C. MuJoCo: MyoBack + exoskeleton

We validated the MyoBack model by conducting trials with a passive exoskeleton. The aim was to compare the forces exerted by the exoskeleton in simulation and experiments for lifting tasks. We modeled the passive exoskeleton, the Auxivo Liftsuit 1.0 (Auxivo, Schwerzenbach, Switzerland) [32], in MuJoCo. A simple model of the passive exoskeleton was implemented respecting the mass distribution and elastic properties of the exoskeleton. For the latter, the two main elastic bands on the back were considered. The elastic bands of the passive exoskeleton were modeled as tendons [33] attached at the level of the 5th thoracic rib and the posterior superior iliac spine (see Figure 3), corresponding to the actual Auxivo elastic band positions. The tendons’ stiffness was calculated to produce an equivalent force to that of the elastic bands. The Auxivo Liftsuit has previously been experimentally benchmarked, where the stiffness of its elastic bands was measured. Nonlinear relationships were identified between the force exerted by the elastic bands and their elongation during lifting and lowering [34]. However, in MuJoCo, only a constant stiffness can be implemented. The nonlinear functions were thus approximated to a unique linear function with stiffness *K*_*Auxivo*_ = *F/*Δ*L* determined to be *K*_*Auxivo*_ = 1527.5*N/m*. Due to the different resting lengths of the elastic band in the Auxivo Liftsuit and the tendons in the back model, the value for *K*_*Auxivo*_ had to be scaled accordingly and the stiffness obtained for the tendons is *K*_*MuJoCo*_ = 526.4*N/m*.

**Fig. 3:**
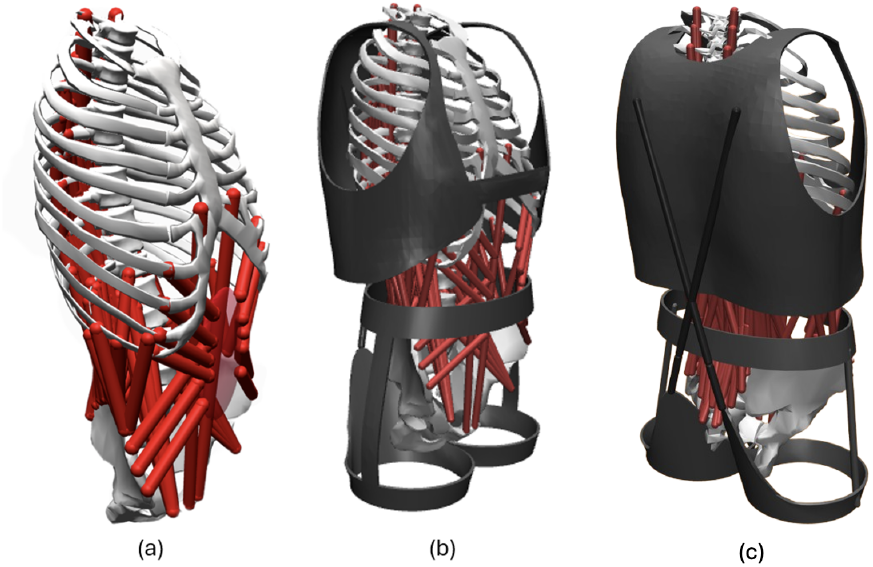
(a) Back model in OpenSim. (b) MyoBack model with exosuit in MuJoCo. (c) Backward view of MyoBack model with exosuit.

The visual meshes of the exoskeleton were built using 3D modeling software separately and then added to the model. The complete MyoBack model is shown in Figure 3. For wider application, a separate version of the MyoBack model is available, with legs incorporated from the MyoLegs model of the MyoSuite framework [35], [36].

## III. EXPERIMENTS

The Auxivo Liftsuit [32] was used to validate the back model. This device, designed to support lifting tasks by reducing strain on the back and hip muscles, helps users handle loads between 5 and 20 kg with less physical stress. Users reported noticeable reductions in muscle strain and an overall improvement in lifting ease while wearing the LiftSuit [37]. Although not unique in the market—other devices like the Darwing Hakobelude (Daiya Industry Co., Ltd., Okayama, Japan) [38] offer similar support—the LiftSuit has been studied in several research contexts. User reports remain promising for its effectiveness, among other settings, in industrial applications [39], [40], [41].

### A. Sensoring the exosuit

First, load cells were integrated into the elastic bands of the Auxivo to measure the actual forces applied by the exosuit onto the user, as shown in Figure 4. One end of the load cell was attached to the metal part of the elastic band and the other to the straps.

**Fig. 4:**
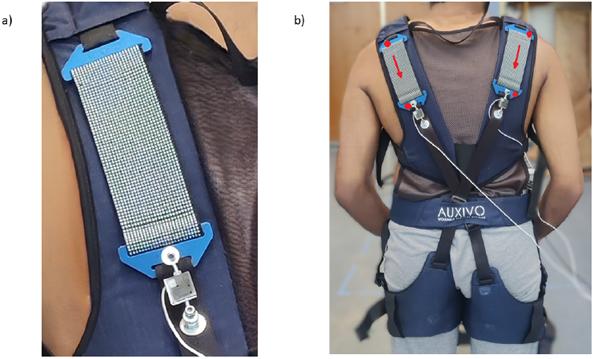
a) Load cell attached in series to one end of the elastic band situated in Auxivo b) Auxivo exosuit with 2 load cells attached to each of the elastic bands. The red arrows on the elastic bands indicate the line of action of stretch force and the red dots indicate the positions of markers at the anchoring points of the elastic bands on the exosuits.

The load cells, placed in series with the elastic bands, measured the deformation of these bands as voltages. These were amplified and digitized before being processed by an Arduino microcontroller. A mapping between the forces exerted and the amplified voltages was known through prior calibration.

### B. Participants

A sample of 9 participants (height in cm: 173.8 *±* 11.1, weight in kg: 73.4 *±* 16.5, age in years: 23.5 *±* 1.4) with no previous experience with exosuits and with no history of back injuries were recruited for the study. All participants provided informed consent. The study was approved by the University of Twente’s ethical review board (Ref. number: 240424).

### C. Apparatus

For this study, data from load cells, reflective markers (Qualisys Medical AB, Gothenburg, Sweden), and IMUs (Movella, Enschede, Netherlands) were used. 63 optical markers were placed on the participant. 56 markers on bony landmarks and body segments and 7 on the exosuit as in [34]. 17 full-body IMU sensors were placed on the subject. Data from IMUs was visualized to examine flexion extension and knee joint angles during static hold trials. Marker data was sampled at 128 Hz and load cell data at 80 Hz. The data from markers was synchronized with the Qualisys Track Manager. This was then manually synchronized with the load cells using a switch during the trials.

### D. Experimental protocol

First, the subjects performed a maximum voluntary contraction (MVC) trial to establish a baseline measurement of each muscle strength. To acquire the MVC for back muscles, the participants were asked to lie on their stomachs and perform reverse crunches against resistance, and vocal motivation was used to ask the participants to contract their muscles as much as possible. Following this, exosuits were worn by the participants and adjusted according to individual user specifications. The participants were asked to perform the following tasks commonly linked with industrial tasks [42].

1) *Static hold trials:* Participants were asked to maintain stoop posture at predefined flexion extension angles of 40°, 60°, and 80° for 3 seconds before returning to an upright posture. Then, participants were asked to perform squats at predefined knee flexion angles of 70°, 90°, and 110°, holding the position for 3 seconds before returning to the initial upright position. Both squatting and stooping movements were repeated three times each. The participants used data from IMUs as visual feedback to maintain the required joint angles.
2) *Dynamic trials*: Participants were instructed to stoop by flexing their trunk to lift an empty box from a table in front of them, then return to an upright posture. They then stooped again to place the box back on the table. Both the squatting and stooping movements were performed at a pace of 40 beats per minute using a metronome, with five repetitions for each movement.

### E. Data analysis

The load cell data was filtered by a zero-lag second-order Butterworth low pass filter of 6Hz. Forces from load cells on both sides of the exosuits were averaged across all 9 participants. To obtain the resultant force acting on the trunk of the user, the measured interaction forces of the right and left elastic bands were summed.

Raw marker data was filtered by a zero-lag second-order Butterworth low pass filter of 6Hz. Marker data was used to compute the stretch forces of elastic bands and joint angles of flexion extension and knee. The previous approach described in [34] for the estimation of forces using markers on the exosuits, as shown in Figure 4, was implemented. In doing so, the change in distance between the markers placed on either end of each elastic band from when the participant was upright was considered band elongation. This value was fed to two polynomial equations representing the loading (flexion) and unloading (extension) phase of a task cycle to estimate forces for each exosuit. The estimated forces from the two elastic bands were summed and averaged across all participants for both exosuits. This conventional marker-based method for determining stretch forces was employed alongside the original sensor-based approach of integrating load cells into an exosuit for verification.

The flexion extension and knee joint angles were computed using marker trajectories in the 2-D sagittal plane. For flexion extension, the angle between the vector joining the markers C7 vertebra and right posterior superior iliac and an arbitrary vector perpendicular to the ground was measured. Similarly, knee flexion angles were computed using the two vectors joining right anterior superior iliac-right lateral femoral epicondyl and right lateral femoral epicondyl-right lateral malleolus.

### F. Simulation

The motion patterns of the experimental trials were replicated in simulation using the MyoBack model with the exoskeleton. For the static hold trials, a sweep was done over the two relevant parameters: flexion extension angle and knee flexion angle. The latter needs to be mapped to the appropriate model joints using marker data to achieve a realistic squat position with the subject leaning forward. For the dynamic trials, marker data was again employed to obtain the joint angles and simulate the complete motion cycle of the stoop/squat.

### A. Static hold trials

In the static stoop results (Fig.5a), the flexion extension angles executed by participants 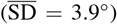 were consistently lower than the predefined values. The MyoBack model with the exoskeleton shows a good reproduction of exoskeleton forces exerted on the human body, notably better with force data derived directly from the load cell sensors (RMSE_lc_ = 4.7N with 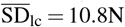) than the marker data (RMSE_m_ = 16.4N with 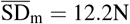). In the case of static squat (Fig.5b), the knee flexion angles match more accurately with the intended ones. However, the uncertainty increases for a deeper squat regarding the knee flexion angle 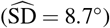 and the measured forces 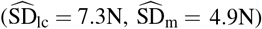. Also, here the simulated exoskeleton forces follow the experimental trend well (RMSE_lc_ = 2.6N, RMSE_m_ = 2.1N).

**Fig. 5:**
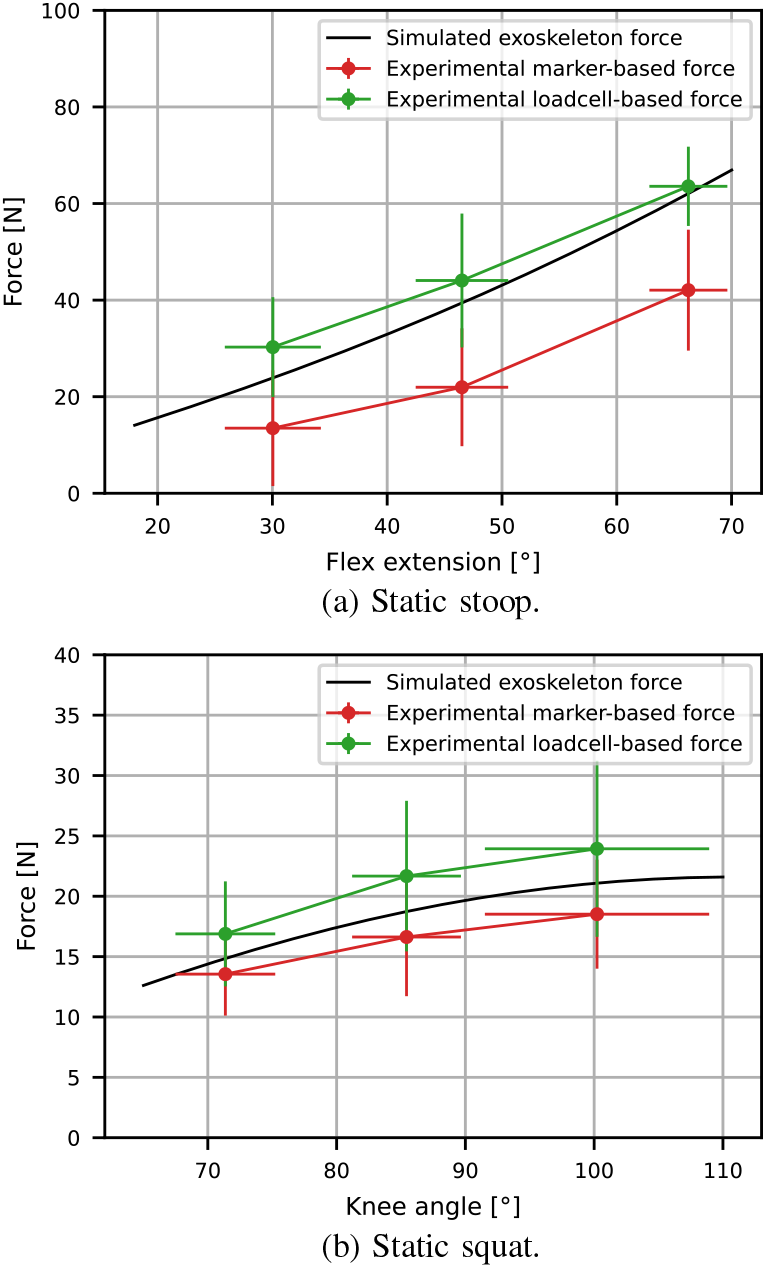
Comparison of experimental static trial exoskeleton force data for stoop and squat motion patterns, derived from load cells and indirectly from markers, with simulation data.

### B. Dynamic trials

To allow for a better comparison, the data was normalized to the percentage of a stoop/squat cycle. For the stoop cycle (Fig.6a), the force data derived from the load cells 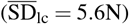 and the markers 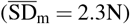 shows similar behavior to the static trials with the latter being consistently lower than the former and the simulated exoskeleton forces matching the former better (RMSE_lc_ = 2.8N, RMSE_m_ = 7.2N). Resembling the static trials, the squat cycle (Fig.6b) shows high uncertainties 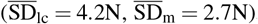. When the squat cycle is simulated using the MyoBack model with the exoskeleton, the reproduced force progressions correlate well with the experimental ones (RMSE_lc_ = 2.9N, RMSE_m_ = 3.8N).

**Fig. 6:**
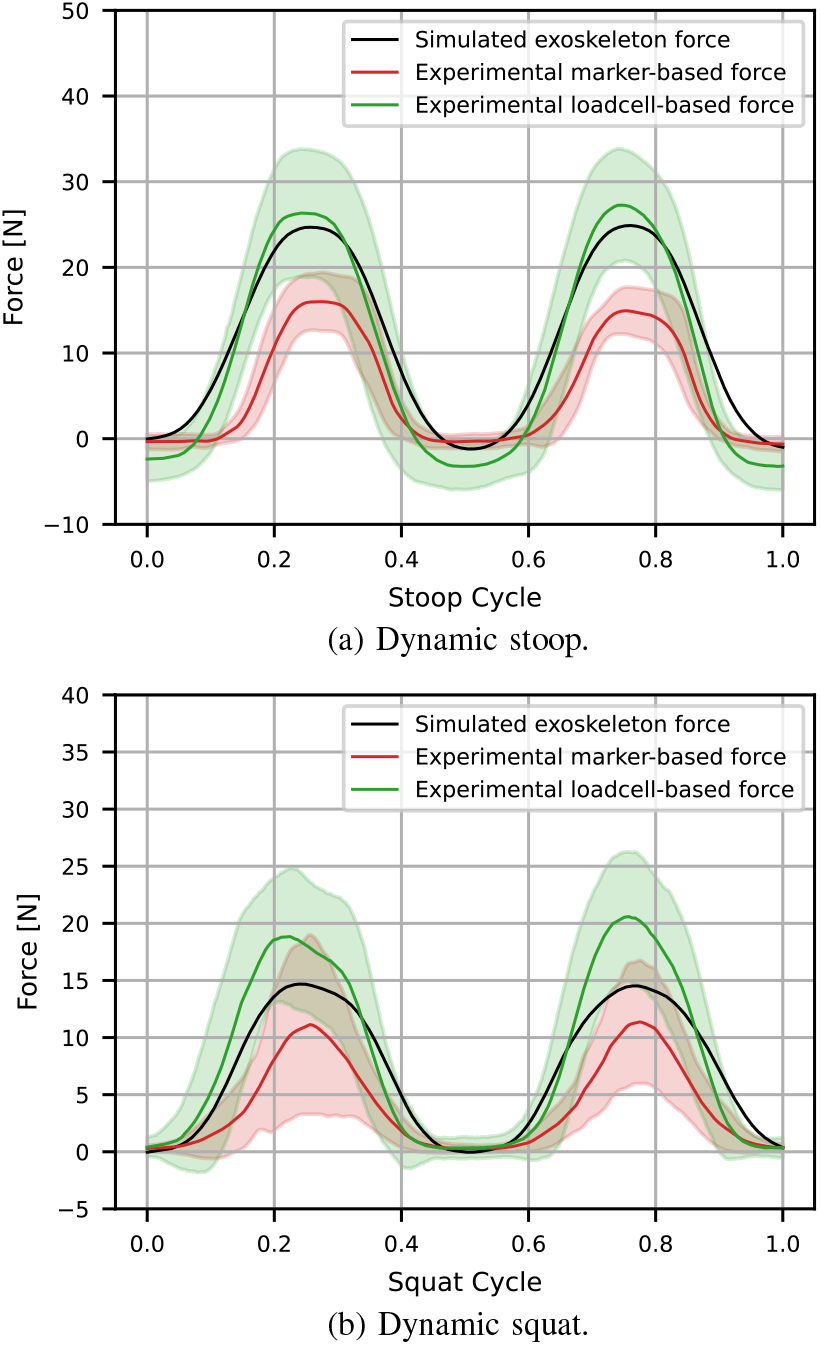
Comparison of experimental dynamic trial exoskeleton force data for stoop and squat motion patterns, derived from load cells and indirectly from markers, with simulation data.

## V. Discussion And Conclusions

This study introduced the MyoBack model, a human back model part of the MyoSuite framework, derived from a physiologically accurate OpenSim model. The two models were compared with respect to kinematics, moment arms exerted by the muscles depending directly on the muscle pathways and the force-length-activation of muscles to ensure the accuracy of the derived model. No discrepancies were observed for the former, suggesting that the geometries and joint relations have been transferred successfully to the MyoBack model. However, regarding the latter two deviations can be seen. This is consistent with OpenSim models of other body parts with similar complexity (number of muscles and DoFs) converted using the MyoConverter, such as the neck model (72 MTUs and 6 joints) [27].

The observed differences stem from the fundamental disparities in how MuJoCo and OpenSim handle physical constraints, such as wrapping surfaces and geometry. OpenSim provides a detailed approach to muscle pathway constraints through specialized wrapping surfaces, while MuJoCo is limited to simple geometries and constraint handling methods. Hence, although MyoConverter facilitated the model conversion, it reached its optimization limit for muscle pathways, leaving some differences unaddressed. Importantly, the direction and trend of the moment arms match well between the two models and the fact that there are no discrete jumps. Also, the muscles themselves are modeled differently in MuJoCo and OpenSim. The possible MTU length may vary for the force-length activation, given that the muscle pathways are different in the two models. But notably, the force exerted at the relative length of the muscle is the same, and the maximum muscle force matches well. This allows for simulation with the MyoBack model with reasonably accurate results.

For empirical validation, the Auxivo Liftsuit, a passive exoskeleton, was modeled in MuJoCo and added to the MyoBack model. Force data derived from load cells and marker information were successfully compared with simulated exoskeleton forces acting on the human body in the context of stoop/squat movements. However, the study’s major weaknesses include a small sample size of nine participants and the use of a standard-sized exosuit, which did not account for individual differences in body size. This led to variable exosuit forces across participants. Additionally, variations in measured flexion-extension angles could be attributed to potential misplacement or slippage of IMU sensors on the lumbar region during repeated or maximum forward bending. Furthermore, the placement of pelvis markers on the exosuit rather than on bony landmarks could have contributed to inaccuracies in computed angles.

Variations in knee flexion angles might also result from offsets in the IMUs placed on the thigh and shank. The significant discrepancies between forces derived from load cells and markers requires further investigation. Notably, load cell data, which directly measures force within the system, is likely to be more reliable. In contrast, markers are prone to slippage over the duration of trials and may be placed inconsistently among participants with varying anatomies.

Future work involves the analysis of EMG data collected from experimental trials, which will enable a more comprehensive validation of muscle activation levels within the MyoBack model. Currently, the validation considers only kinematics and muscle pathways. Still, exploring whether a reinforcement learning (RL) agent trained to perform specific static and dynamic tasks will exhibit activation patterns similar to those observed in human trials will be informative. The MyoBack model has the potential to extend its applications to various aspects of human locomotion and agility, including object lifting, balancing, and running, which remain unexplored in the current study. Additionally, integrating this model with both active and passive exoskeletons across the full body could further be used for rehabilitation and industrial design purposes and benefit populations such as the elderly and workers.

